# Relatedness coefficients and their applications for triplets and quartets of genetic markers

**DOI:** 10.1101/2022.10.28.514212

**Authors:** Kermit Ritland

**Affiliations:** Biodiversity Research Center, University of British Columbia

## Abstract

Relatedness coefficients which seek the identity-by-descent of genetic markers are described. The markers are in groups of two, three or four, and if four, can consist of two pairs. It is essential to use cumulants (not moments) for four-marker-gene probabilities, as the covariance of homozygosity, used in 4-marker applications, can only be described with cumulants. A covariance of homozygosity between pairs of markers arises when populations follow a mixture distribution. Also, the probability of four markers all identical-by-descent equals the normalized fourth cumulant. In this paper, a “genetic marker” generally represents either a gene locus or an allele at a locus.

Applications of three marker coefficients mainly involve conditional regression, and applications of four marker coefficients can involve identity disequilibrium. Estimation of relatedness using genetic marker data is discussed. However, three- and four-marker estimators suffer from statistical and numerical problems, including higher statistical variance, complexity of estimation formula, and singularity at some intermediate allele frequencies.

## INTRODUCTION

Relatedness is a general term for the level of genetic similarity between individuals, and is measured by the sharing alleles identical-by-descent (MalÉcot 1948; Pamilo and Crozier 1982). Relatedness is quantified with gene identity coefficients, which characterize both the pattern and the frequency of identity-by-descent. The unit of observation is normally a pair of marker loci, and the object of estimation is the kinship coefficient or the coefficient of relationship. In this paper, the unit of observation is extended to triplets and quartets of genes, allowing the opportunity to characterize additional parameters of population structure.

Relatedness may be estimated with genetic markers (Morton et al. 1971) and for pairs of marker genes, many computer programs are available for estimation of relatedness (Wang 2014), in particular for “pairwise relationship”, such as the *r* of Queller and Goodnight (1989). However, the equations for pairwise relationship are not extendable to three or four genes, as the covariances and higher moments need to be defined in new ways.

With more than two markers the situation becomes much more complex. Estimators for three and four marker-gene measures of relatedness have recently been proposed. SAMANTA (2009) provided the first estimator for three genes. Ackerman *et al*. (2017) examined all four genes and described the estimation of seven of eight coefficients of relatedness. Multiallelic data have information about all eight coefficients, but they used a biallelic model which provides just seven degrees of freedom, constraining their space of estimates. In addition, they did not use cumulants and their moments are normalized differently than will be here.

Cumulants are of use in certain problems in quantitative genetics (Burger 1991; Turelli and Barton 1994) and compared to cumulants have more useful theoretical properties (Kendall *et al.* 1977), but ordinary moments are sufficient for two and three-gene fixation indices. Ritland (1987) found cumulants instead of moments were an essential component of four-gene fixation indices. Fourth order cumulants are needed to specify the probability of gene identity for all four genes, and also to describe identity disequilibrium as the “covariance of covariances”. As an example of the necessity of cumulants, for population gene frequency *p*, the fourth central moment for four genes, denoted (*X*_1_,*X*_2_,*X*_3_ and *X*_4_), is 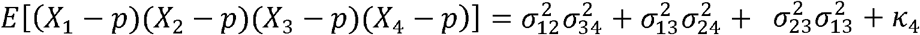 where 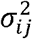 is the covariance of *X*_*i*_ and *X*_*j*_ and *κ*_4_ is a fourth-order cumulant which does not appear in moment-based treatments.

The probabilities developed in the paper are used to estimate relatedness in a population with polymorphic genetic markers. Any numbers of alleles at a single locus are allowed in the equations, but practically, one has two and at most three alleles with SNP types of data, and for tractability, we use a three-allele model to examine a special situation with four genes. These models and estimation procedures are readily applicable to the emerging mountains of genome data.

## APPLICATIONS OF MARKERS

### Definitions

Between pairs of genetic markers, the *coefficient of relationship* measures the degree of consanguinity (e.g., the probability that markers are identical by descent, termed ibd). The coefficient relationship equals twice the *kinship coefficient*. The *inbreeding coefficient* is the probability that a pair of markers within one individual are ibd. With more than two markers, the coefficient of relationship is more broadly defined with groups of markers (two, three or four). With four markers, there are nine *modes of Jacquard’s gene identity*, with ibd genes connected by lines. The *normalized central moment* gives the probability of ibd of all markers. At the level of four markers, *cumulants* are necessary to describe *identity disequilibrium* (the excess of identity between marker pairs). The covariance of cumulants forms the machinery of higher order interactions.

### Relatedness and two markers

The two-marker coefficient of relatedness is used for many inferences with genetic markers, mainly involving pairs of genes sampled between two individuals (“ coefficient of relationship”or pairs of genes sampled within one individual (the “inbreeding coefficient”). Analysis of data with two marker measures are ubiquitous (Wang 2014) and the two-marker probabilities are often incorporated into probabilities of groups of three and four genes.

### Regression and three markers

The three-marker relationship coefficient is the probability that the three marker loci have alleles all identical-by-descent (Figure 1a). This coefficient, *G*, is usually combined with two-allele coefficients for biological interpretable parameters, useful at least for problems involving mating systems or kin selection. In the theory of mating system estimation, the “effective selfing rate” is the genetically equivalent rate of selfing caused by all types of biparental inbreeding (Ritland 1985); the effective selfing rate of individual *A* equals 2*R*-*G*, where *R* is the relatedness between mates and G the third moment involving the two maternal and single paternal allele (Figure 1b). In the theory of kin selection, the regression coefficient of relatedness is used and, properly, a three-gene model is needed (Figure 1c), as shown by Michod and Hamilton (1980), where their Equation 18 depends upon whether the reference genotype is homozygous (18a) vs. heterozygous (18b). Note that both the effective selfing rate and the regression coefficients of relationship can be asymmetrical when inbreeding coefficients differ between the two relatives.

**Figure 1.**
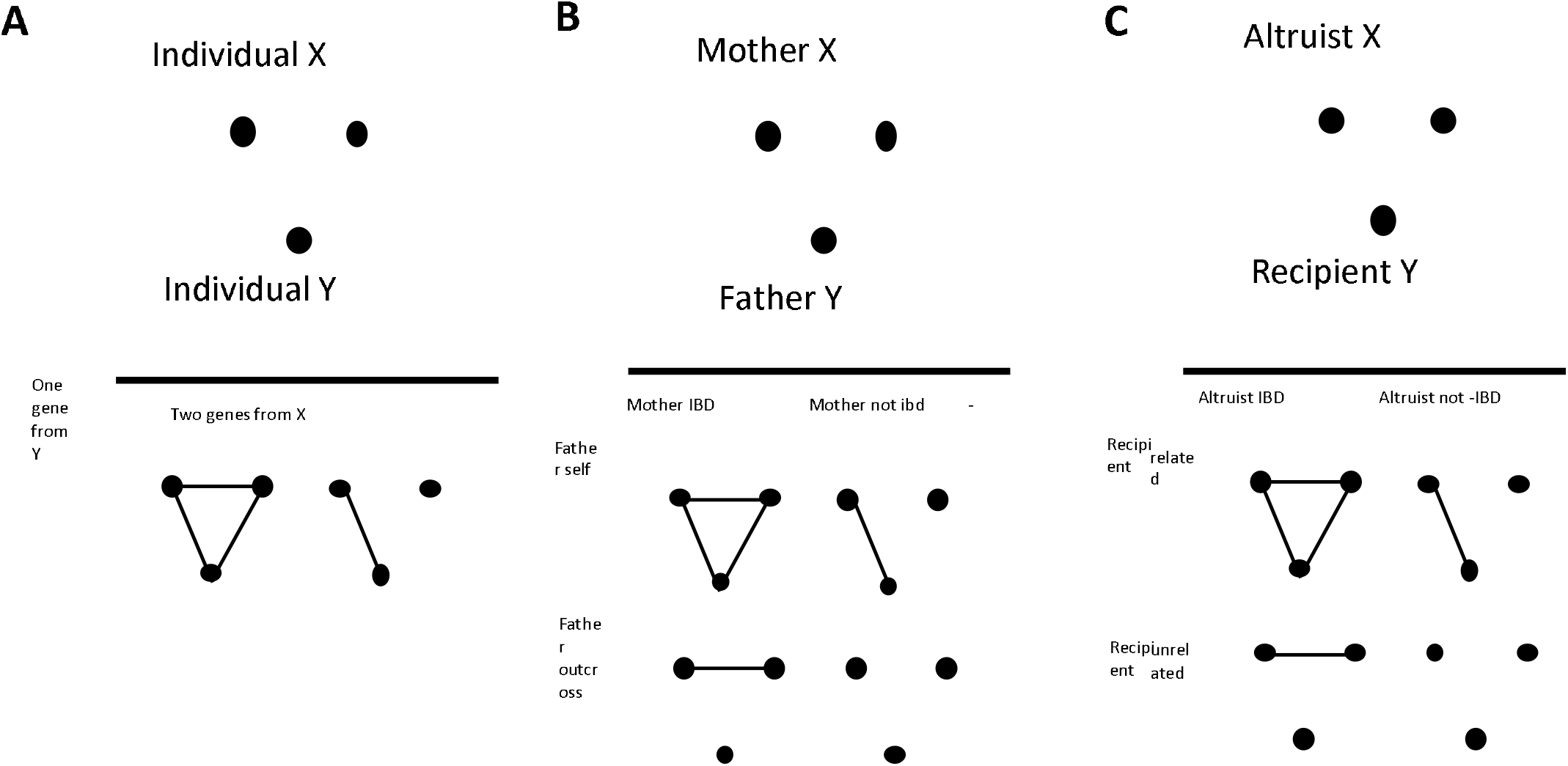
Three cases where three-gene modes of gene identity are used. (A) Effective selfing model involves two genes of one parent, (B) progeny pair model, (C) Altruist-recipient. Identical genes are linked by lines. In the effective selfing model, two of the genes are from the maternal parent and the other gene is the paternal contribution.

Identity disequilibrium and four markers

Between two diploid individuals, there are 15 patterns of gene identity (Liu 2005). A pair of individuals can share two, three or four markers, and at each level, allelic similarities can describe aspects of relatedness. After Jacquard (1966); Jacquard (2012), for four marker genes and two individuals, there are nine condensed identity modes, denoted as Δ_*i*_ (Figure 2a). There are eight independent parameters of relatedness: three pairwise measures (*F*_*A*_, *F*_*B*_, *R*), two three-way measures (*G*_*A*_, *G*_*B*_), and three four-marker measures (*F*_*AB*_, *R*_*AB*_, *H*). At the highest level, the measures are much different as the four-marker measures *F*_*AB*_, *R*_*AB*_ are covariances (not identities). The four-marker parameter, *H*, is the probability that four markers are identical. As well, the quantities must be defined as cumulants. Cumulants equal moments up to order three, but fourth-order moments do not equal fourth order cumulants.

**Figure 2.**
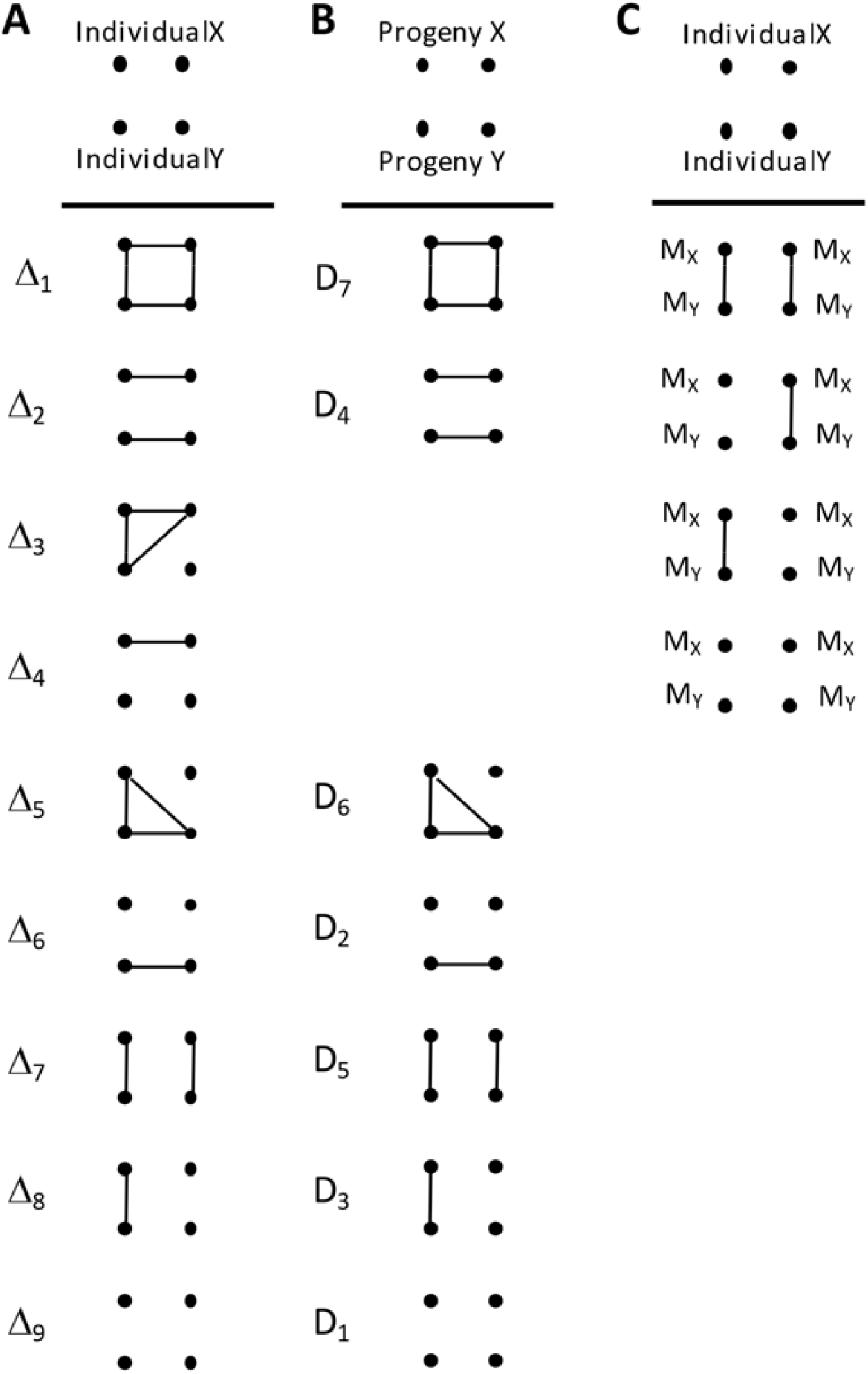
Two cases where four-gene modes of identity are used. (A) The general case where each of the nine identity modes are inferred (B) A model to fit progeny pairs to mating system parameters (C) The inference of heritability in the field, where M is the marker and Q the quantitative trait.

The fourth central moment equals κ_4_ +*F*_*AB*_ +2 *R*_*AB*_ (note the covariances between second moments enter this expression) and the normalized cumulant κ_4_ (equivalent to *H*) equals the probability of identity of all four marker genes. While the variance is a measure of the spread of the distribution, kurtosis is a measure of the “peakedness” of the distribution of random variables, and infrequent extreme deviations contributing excessively to this statistic (de la Rosa and Moreno muÑoz 2008).

While applications of three and four gene measures are in their infancy, at least, the skew and kurtosis as measured by higher moments can help remove bias in DNA forensics caused by genotyping error (Weir 1994).

In the four-marker model, many possibilities exist about attaching meaning to each of the Δ_*i*_. One example is the progeny-pair model (Ritland and Leblanc 2004) where *A* and *B* are two progeny of the same mother plant (Figure 2b). At another more abstract level, two of the genes can be markers and two are quantitative trait loci (Figure 2c). If identity disequilibrium is present, the regression of phenotypic similarity (QTL) on estimated relationship (markers) gives an estimate of heritability “in the field” (Ritland 2000).

### TWO, THREE AND FOUR MARKER PROBABILITIES

At any level of comparison, associations are measured as the frequency of a given configuration (allele “state”) divided by the denominators in Table 1. These denominators are termed “normalization constants”, and are the maximum possible value of the numerator. Some of these normalized measures of association arise naturally in the derivations below.

**Table 1.** Examples of the statistical variances of relationship coefficients when estimates are based upon a single locus and when the true values are zero (“x”denotes not estimable). See equations 3 and 6 for definitions of G and H.

| 395 | Array of $p$ | $F$ | $G_A$ | $G_B$ | $\phi_{XY}$ | $H$ |
| --- | --- | --- | --- | --- | --- | --- |
| 396 | 0.6,0.3,0.1 | 2.93 | 73.5 | 74.3 | 12.96 | 7.63 |
| 397 | 0.5,0.3,0.2 | 2.32 | x | x | 11.66 | 3.66 |
| 398 | 0.4,0.3,0.2,0.1 | 1.58 | 0.19 | 7.79 | 13.05 | 2.18 |

#### 1. Probabilities of two-marker relationship

From Equation (7) of Ritland (1987), which follows Kendall *et al*. (1977 eq. 13.36), the frequency of gametes with allele *i* and with allele *j* is

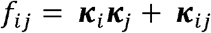

The two-marker coefficient of relationship can be estimated from the frequencies of each allele in a sample. For any given allele, say *A*_*i*_, it derived by equating the observed frequency of homozygotes to that expected by the above equation

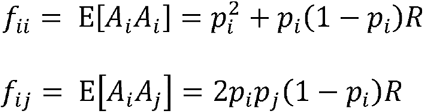

The likelihood of the data, given *R*, 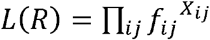. Solving for *R* gives estimators based upon pairs of alleles *A*,

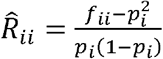

The estimate of *R* for allele *i*, estimates are combined across alleles as

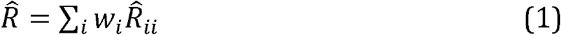

where the weights *w*_*i*_ sum to unity.

These weights are found by finding the *w*_*i*_ that minimize **w**^T^**Vw**, where **w** is an *n* element vector of weights, and **V** the *n*x*n* variance-covariance matrix of allele-specific estimates (for details see Ritland (1996)). The weights require prior specification of true relatedness. With zero prior *R*, the weight for allele *A*_*i*_ is 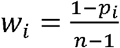. An *m*-allele locus receives the weight (*n*_*m*_-1), giving the estimator for *r* given by equation 5 in Ritland (1996).

#### Probabilities of three-marker relationship

The three-marker relationship coefficient is the probability that three sampled marker genes are all identical-by-descent. From Equation (7) of Ritland (1987), which follows Kendall *et al.* (1977 eq. 13.36), The joint frequency of markers *i, j* and *k* is

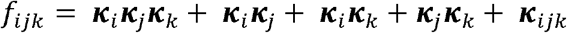

This written in conventional population genetic terms as

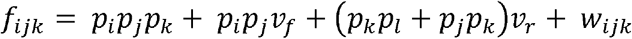

Where alleles *i* and *j* are from one individual and allele *k* from a second individual. The cumulants are written in bold face to emphasize they have a random component that may covary.

From Equation (7) of Ritland (1987), there are three primary patterns

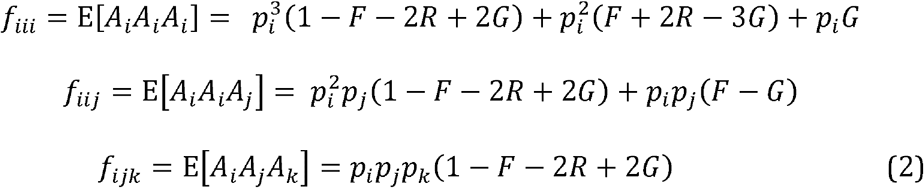

Where the order is irrelevant (*A*_*i*_ *A*_*i*_ *A*_*j*_, *A*_*j*_ *A*_*i*_ *A*_*j*_ and *A*_*i*_ *A*_*j*_ *A*_*j*_ are equivalent). The genotype frequencies are mixtures of marker gene identity: *G* is the probability that all three markers are ibd, *R-G* is the probability of ibd of one pair of markers, *F*+2*R* -3*G* for two pairs, and and 1-F-2*R*+2*G* is the probability of no ibd among the three markers).

Solving for *G* in Equation (7) gives three probabilities involving *G*,

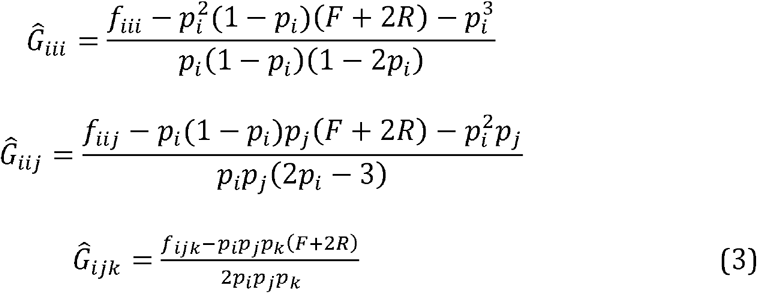

*G* is a normalized third central moment and the normalization constant depends upon the pattern of subscript.

Each allele can provide an estimate of *G*, denoted 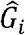, and its weighted estimate across possible alleles *i* is

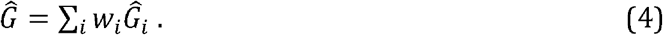

This represents a “linear estimator” of *G*. The weights are derived in the appendix. The best alternative to linear estimation is maximum likelihood.

### Probabilities of four-marker relationship

From Equation (7) of Ritland (1987), which follows Kendall *et al*. (1977 eq. 13.36),

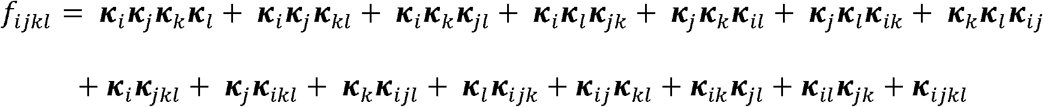

The cumulants ***k***_*i*_ are similar to moments and covariances and may have a random component that may covary with other cumulants. The subscripts indicate alleles. The recursion equation is

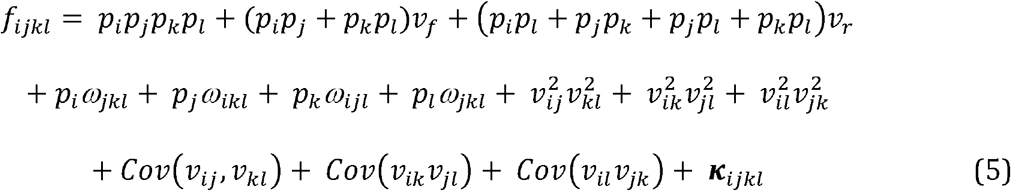

Where the *v* terms are second order covariances. When there is a mixture model (which creates the covariances), each subpopulation m, contributes to the mean cumulant across pooled m. The term *k*_*i,m*_*k*_*j,m*_*k*_*k,m*_*k*_*l,m*_ contributes to all 18 population level moments, the term *k*_*i,m*_*k*_*j,m*_*k*_*kl,m*_ contributes to six population level moments, and so on. However, the quantitative extent of these contributions are complex and beyond treatment here. Regardless, that subpopulation cumulants “distill” to the same assortment of cumulants, albeit with perhaps slightly different values. where the covariance terms are across the mixture terms *m*. This is a finite mixture model, needed when a single component distribution is inadequate (Mclachlan *et al.* 2019). These can get complex but Withers *et al.* (2015) does provide the first known expressions for cumulants used available computer technology (an equation solver), not available in 1987 to KR. Cumulants are allowed to vary across the mixture, and that this results in effective covariance between second-order cumulants. The expectations taken across *m* result in changes to the above expression due to associations among the *ijkl* across *m*, that causes the cumulants to be associated in a certain way, since for example, *E*[*k*_*i,m*_*k*_*j,m*_] ≠*k*_*i*_*k*_*j*_.

The associations between *pairs* of markers is termed identity disequilibrium. If one pair of alleles is heterozygous, it is more likely the second pair is also heterozygous. This is a four-gene marker measure that has been neglected due to inordinate attention to linkage disequilibrium. Identity disequilibrium has classically been characterized as the excess of homozygosity above that expected from the squared gene frequencies (Hill 1975) (Ohta 1980). The identity excess is closely correlated to the expectation of the total squared linkage disequilibrium (Takahata 1982). Some of the problem is that haploid gametes are not directly assayed but rather imputed (Vitalis and Couvet 2001b).

We can add a cumulant to the equation for the probability of identity-by-state. From equation 3.78 in Kendall *et al.* (1977), the moments about the mean for two squared random variables (the alleles present at each locus) equals

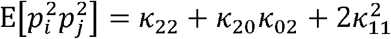

Whose form corresponds to 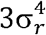 for the fourth central moment with the difference that a the cumulant *k*_22_ is added. Vitalis and Couvet (2001a) and others have given estimator for identity disequilibrium which omits this cumulant.

The four-allele case introduces higher-order associations and brings with it new statistical problems. Among four alleles, two new measures arise. The first is termed *H* and is the probability that all four alleles are identical-by-descent. The other two have not been recognized in the literature, perhaps because they invoke the existence of cumulants, which differ from the corresponding moments with products of four or more variates.

The first, termed *R*_*AB*_, is the probability that both alleles in the first relatives are identical-by-descent to both alleles in the second relative. The second, termed *F*_*AB*_, is the probability that both individuals have both marker genes identical-by-descent. Thus, the three unique four-allele measures are

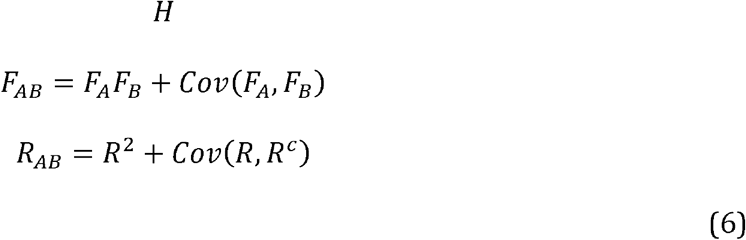

the covariances between second moments, *Cov*(*FA,FB*) and *Cov*(*RAB, R*′_*A*_*B*′), exist only when the distribution of gene frequency follows a mixture distribution where subpopulations vary for *F* and *R*.

We can rewrite Equation (5) as

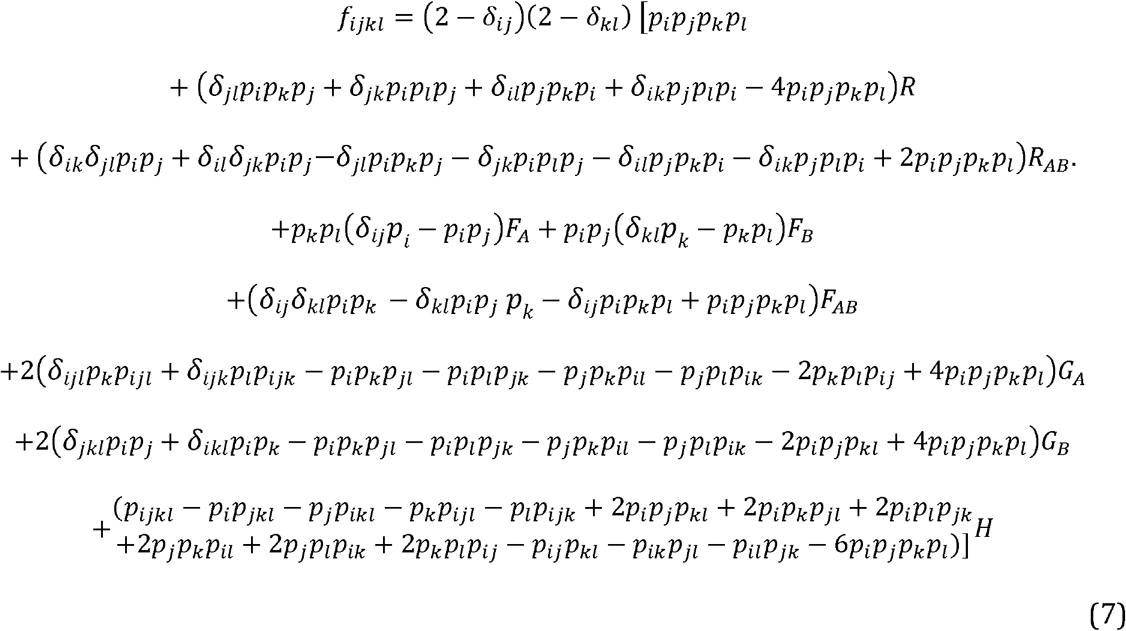

where, for shorthand, *p*_*ij*_ = *δ*_*ij*_ *p*_*i*_.

In this expression, there are eight relationship coefficients (*R*_*AB*_, *R*_*ABAB*_, *F*_*A*_, *F*_*B*_, *F*_*AB*_, *G*_*A*_,*G*_*B*_,*H*), which in principle will specify eight different classes of marker genotypes. This probability of four alleles,, *f*_*ijkl*_, is then fitted to the observed frequencies in a sample.

For equations that solve for all eight parameters, the choice is somewhat arbitrary but a natural set of eight classes, in which identity-by-state mirrors the identity-by-descent, is: *A*_*i*_*A*_*i*_*A*_*i*_*A*_i_ (all identical by state, or “ ibs”), *A*_*i*_*A*_*k*_*A*_*i*_*A*_*k*_, *A*_*i*_*A*_*i*_*A*_*k*_*A*_*k*_, (two pairs ibs), *A*_*i*_*A*_*i*_*A*_*i*_*A*_*k*_, *A*_*i*_*A*_*k*_*A*_*k*_*A*_*k*_, (one triplet ibs) and *A*_*i*_*A*_*i*_*A*_*j*_*A*_*k*_, *A*_*i*_*A*_*j*_*A*_*k*_*A*_*k*_ and *A*_*i*_*A*_*j*_*A*_*j*_*A*_*k*_. (one pair ibs between A and B). Thus we seek the expected frequencies in the vector (*f*_*iiii*_, *f*_*ijij*_, *f*_*iijj*_, *f*_*iiij*_, *f*_*iijk*_, *f*_*ijik*_).

The frequency of *A*_*i*_*A*_*i*_*A*_*i*_*A*_*i*_, is obtained from Eq (7) where all δ=1 and all marker frequencies are *p*_*i*_:

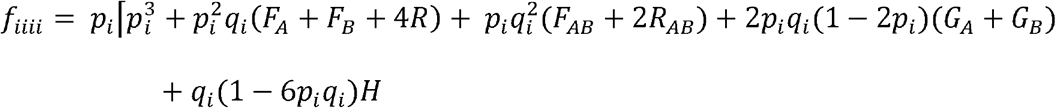

Likewise, the frequency that A and B are both heterozygous for *A*_i_ and *A*_j_ is, irrespective of order or phase,

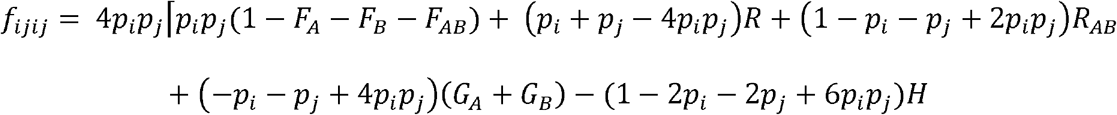

and homozygous for alternative alleles *A*_*i*_ and *A*_*j*_ is

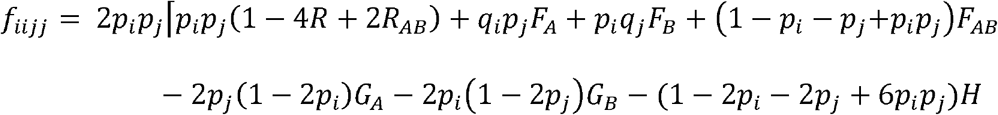

for triplets of identity-by-state *f*_*iiij*_

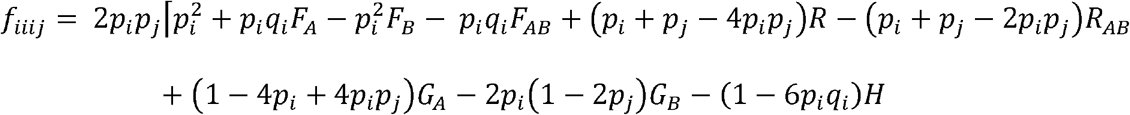

Finally, a single allele pair *ii* can be shared only within individual *A*,

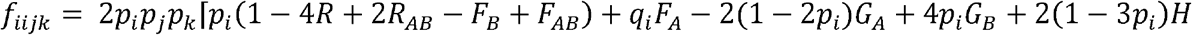

or shared only once between *A* and *B*:

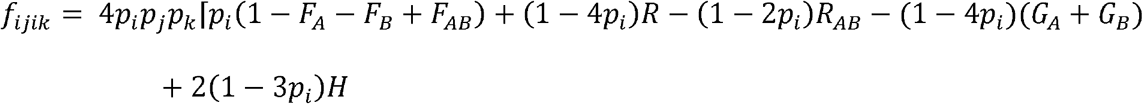

The expressions for *f*_*ijjj*_ and *f*_*ijkk*_ are obtained by symmetry, and the expression for *f*_*jjk*_ is summed over all four pairings of j between A and B: *f*_*ijjk*_ *f*_*ijkj*_ *f*_*jijk*_ and *f*_*jikj*_.

The appendix gives the 8×8 matrix of probabilities of observing the marker gene frequencies f given the relatedness coefficients. Of course, this depends upon the particular array of *f* ’s used (there are others than the above). In this case, the determinant of the matrix is 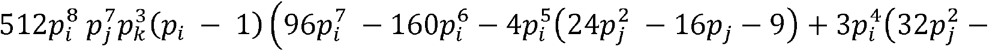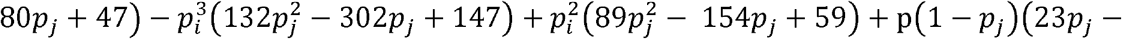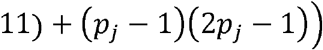. That it is non-zero indicates all 8 parameters are jointly estimable, but a linear approach which uses residuals to simplify things is needed at this point.

### Joint values of H and identity disequilibrium

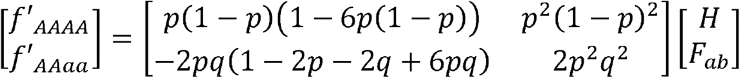

whose solution is

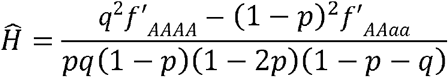

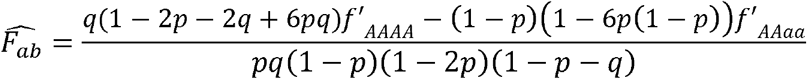

Joint estimates of H and joint identity disequilibrium

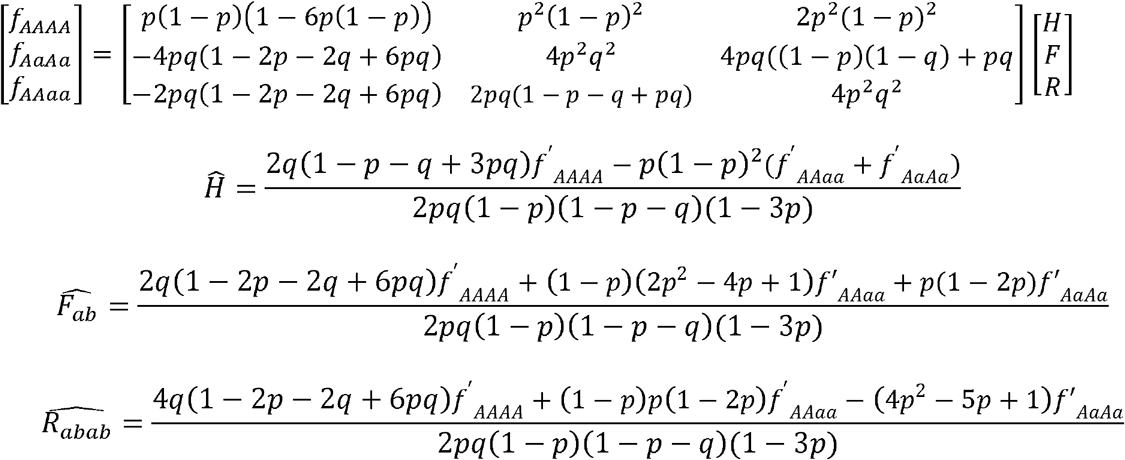

The denominator shows that at least three alleles required in the population, and marker frequencies of *p*_*i*_ =1/3 are non-informative.

Identity disequilibrium has classically been characterized as the excess of homozygosity above that expected from the squared gene frequencies, as proposed by Hill (1975) and Ohta (1980). The identity excess is closely correlated to the expectation of the total squared linkage disequilibrium (Takahata 1982). Some of the problem is that haploid gametes are not directly assayed but rather imputed (Vitalis and Couvet 2001b).

The “classic” procedure for estimating identity disequilibrium involves comparing observed vs. expected double homozygotes. From Vitalis and Couvet (2001a), one such estimator can be written in the form,

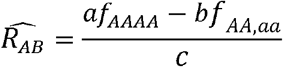

where *a* = 6(6*pq* − 2*p* − 2*q* + 1), *b* = (1 − 6*pq*), *c* = 6*pq*(1 − *p* − *q*)(1 − 3*p*). The numerator is positive as *a* is always larger than *b*, but denominator can be either positive or negative and negative values occurs when *b* is larger an *a*. Such is life with four genes. It is also amazing the four-gene frequencies *f*_*AAAA*_ and *f*_*AA,aa*_ are directly used in an effective two gene estimator, as *f*_*AAAA*_ is the observed double homozygotes and *f*_*AA,aa*_ represents homozygotes expected with no zygotic association. Equally amazing is that Vitalis and Couvet (2001a) and others have given estimators for identity disequilibrium which omits this cumulant.

## DISCUSSION AND CONCLUSION

A main feature of higher-order relatedness is the covariance of homozygosity between pairs of marker genes, this is effectively a covariance of second moments. Such a “covariance of covariance” arises when pedigrees occur in a mixture distribution (Mclachlan *et al*. 2019). Such a distribution generates the genomic variation of homozygosity necessary for the existence of covariance of homozygosity between individuals at specific loci. The simplest mixture distribution is that of two populations with gene frequency *p*+*a* and *p*-*a*; in this case covariance of heterozygosity, after mixing in equal proportions, equals *a*^4^+6*a*^2^*p*^2^.

Another feature of higher-order relatedness is that the four-marker coefficient of gene identity must be described with cumulants and not moments. As an example of the necessity of cumulants, for population gene frequency p, the fourth central moment for four markers, denoted (*X*_1_, *X*_2_ *X*_3_, and *X*_4_), is 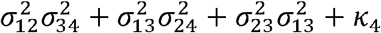 where 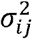 is the covariance of *X*_*i*_ and *X*_*j*_ and *k*_4_ is a fourth-order cumulant which does not appear in the moment. Some type of term (not involving the product of variances) is needed for *k*_4_ and it could be any rational number. In summary, incorporating cumulants into four marker measures only requires some value *X* in the expansion of the fourth central moment 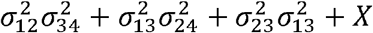 and this *X* is numerically estimated in the same way as the lower order cumulant terms.

Cumulants do have useful properties for models of quantitative traits, the most important is that the cumulant of the sum of two random variables X+Y is M(X+Y)=M(X)+M(Y); differential equations for models of selection on quantitative traits that involve cumulants are simpler than models involving moments (Burger 1991; Turelli and Barton 1994). This cumulant will also be key in deriving a marker-based estimator for *Q*_*st*_ (Ritland in prep) and for a portrayal of higher order population structure that separately accounts for both the correlation of relationship and the squared linkage disequilibrium (Ritland in prep)

We give probabilities of relationship for a homogenous population of just one generation. Such populations are most commonly assayed in genomics, however it should be noted that the levels of nucleotide variation (for SNPs) is not high and loci with more than two alleles are uncommon; in fact, only about 5% of human SNPs are triallelic (Cao *et al.* 2015) although microsatellites and other types of repeat markers show greater variation. Reconstruction of pedigree relationship has traditionally involved cumbersome graph-tracing algorithms, and simpler recursive methods which require at least two generations of records (Karigl 1981; Thompson 1988; Whittemore and Halpern 1994). Also, current recursive methods (Zheng *et al.* 2018) assume a known pedigree (Kirkpatrick *et al.* 2018). This is somewhat like estimating *Q*_*st*_ with current methods, where aspects of the pedigree must be known.

### Normalization constants

Relatedness coefficients are obtained by calculating the pairwise covariance of relatives and dividing it by a normalization constant that converts the covariance into a correlation. This constant is the maximum possible value that the covariance can take. For cases where pairwise comparisons involve the frequency of identical genotypes, it is simple to calculate as a binomial variance. For the two-marker relationship coefficient as described in Equation (1), the maximum covariance between two genotypes, conditioned upon observing allele *i*, is E[*A*_*i*_ *A*_*i*_] E[*A*_*i*_]^2^ = *p*_*i*_(1-*p*_*i*_) when R=1. Likewise, the three marker coefficient has a normalization constant of *p*_*i*_(1-*p*_*i*_)(1-2*p*_*i*_) for *A*_*i*_ *A*_*i*_ *A*_*i*_., and the four-marker coefficients are normalized by *p*_*i*_(1-*p*_*i*_)(1-6*p*_*i*_(1-*p*_*i*_)) for *A*_*i*_ *A*_*i*_ *A*_*i*_ *A*_*i*_. The normalization constants for combinations of alleles falls out of the analyses.

Ackerman *et al.* (2017) provided a different set of normalization constants for three and four marker measures than given here. In their Equation 6, they normalized the third central moment by the geometric mean gene frequency of the three central moments (rather than by *p*(1-*p*)(1-2*p*) as done here). Their justification was a similarity of this “third moment correlation” to a bivariate correlation formula. Their normalization constant for the four-marker coefficient (Equation 8) involves a parameter α that mixes the unknown proportions of the two types of higher order identity (all identical vs. 2 pairs identical), resulting in an inference that may be subject to biases.

### Estimation

Table 2 gives example estimates of *R, G* and *H* at a single locus. As expected, the variances decrease with numbers of alleles. While the variances for *R* are reasonable, those for *H* and the variances for the identity disequilibrium *R*_*AB*_ are quite large. Interestingly, and not remarkably, the variances for G are all over the place and its is not even estimate in one case (p=0.5).

**Table 2.**
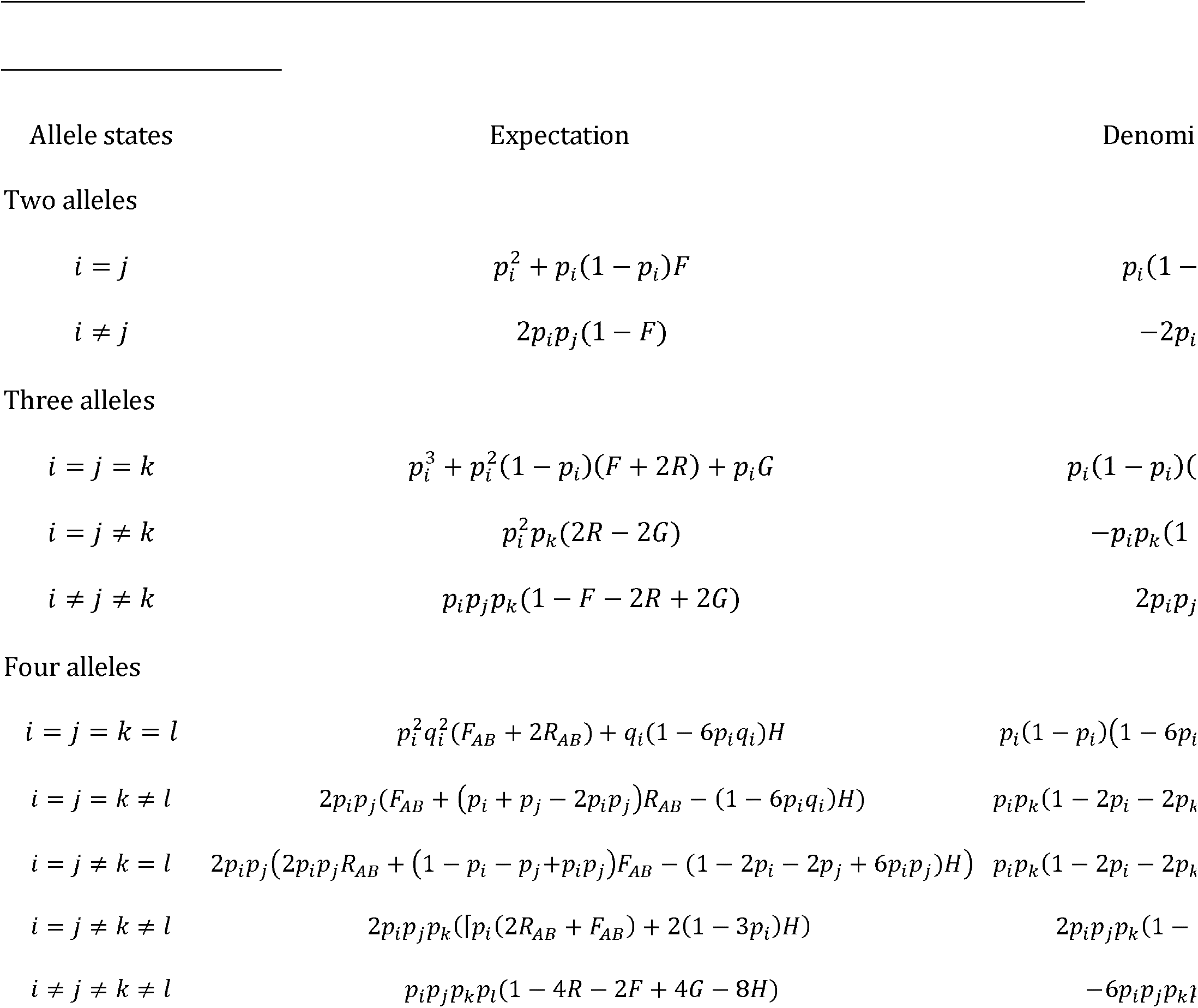
Allele states and denominators of cumulants.

Likelihood requires numerical solutions which introduces complications, as the numerical solution is normally iterated until convergence. We used the Expectation-Maximization method which has slow convergence. When using likelihood, I found the number of marker loci needed for adequate convergence was about 20 loci for three-gene coefficients and roughly double at 30-50 loci for four gene coefficients. Interestingly, it was found that loci with fewer alleles are more likely to give convergent estimates because the problem with non-convergence arises when relatives do not share the same marker allele.

Calculating the probabilities of higher-order relationship poses an interesting set of obstacles. Equation solvers such as Derive or Mathematica help with the derivation and interpretation of complicated formulae. The coefficients can also undefined at certain intermediate frequencies (*p*=1/2 or 1/3) and show high statistical variance about those frequencies (Ritland 1987).

#### Possible future approaches

The complications of correctly estimating population structure are discussed and treated by Weir and Goudet (2017). They developed two-marker moment estimators that can describe the “relativity” and this requires an explicit reference population. They develop their estimator in a multilevel approach (within individuals, between individual within populations, and between populations) which promoted a unified treatment of relatedness and population structure. Clearly, further progress will depend upon adequate definitions and applications of models.

“Relatedness mapping” (Albrechtsen *et al.* 2009) uses relatedness to identifying causative mutations, using the principle that affected individuals share higher relatedness about the mutation. A somewhat related activity is “IBD mapping”, in which segments of identity-by-descent (IBD) present in high-density genomic data are used to map casual variants (Browning and Browning 2012). However, the data by itself only reveals the presence of the variant.

Other fields have adopted the use of cumulants, which may show new approaches that population genetics can undertake. The central 4^th^ cumulant has been used to detect early stages of termite infestation, as it can separate termite alarm signals from background noise (de la Rosa and Moreno MuÑoz 2008). Advances in electrophysiological and imaging techniques are used to study the synchrony of neuron cell firings in the brain (Staude and Rotter 2010), and have highlighted the need for correlation measures that go beyond simple pairwise analyses, taking advantage of the “interaction property” of higher order cumulants as measures of correlation (Staude *et al.* 2010). In information systems, a “covariance of covariance” approach for individual pixels has been developed for image description and classification (Serra *et al.* 2014).

## ACKNOWLEDGMENTS

Joe Felsenstein provided many helpful comments on my first and only sabbatical. This work was supported by NSERC grants to KR

